# REST is a major negative regulator of endocrine differentiation during pancreas organogenesis

**DOI:** 10.1101/2021.03.17.435876

**Authors:** Meritxell Rovira, Goutham Atla, Miguel Angel Maestro, Vane Grau, Javier García-Hurtado, Maria Maqueda, Jose Luis Mosquera, Julie Kerr-Conte, Francois Pattou, Jorge Ferrer

## Abstract

Understanding genomic regulatory mechanisms of pancreas differentiation is relevant to the pathophysiology of diabetes mellitus, and to the development of replacement therapies. Numerous transcription factors promote β cell differentiation, although less is known about negative regulators. Earlier epigenomic studies suggested that the transcriptional repressor REST could be a suppressor of endocrine gene programs in the embryonic pancreas. However, pancreatic *Rest* knock-out mice failed to show increased numbers of endocrine cells, suggesting that REST is not a major regulator of endocrine differentiation. Using a different conditional allele that enables profound REST inactivation, we now observe a marked increase in the formation of pancreatic endocrine cells. REST inhibition also promoted endocrinogenesis in zebrafish and mouse early postnatal ducts, and induced β-cell specific genes in human adult duct-derived organoids. Finally, we define REST genomic programs that suppress pancreatic endocrine differentiation. These results establish a crucial role of REST as a negative regulator of pancreatic endocrine differentiation.

## INTRODUCTION

Progress in our understanding of the transcriptional mechanisms underlying pancreatic β cell differentiation has been crucial for recent advances in the development of regenerative therapy strategies for type 1 diabetes mellitus, including efforts to generate functional β cells from stem cells, organoids or *in vivo* reprograming (Rezania *et al*., 2014; Zhou and Melton, 2018; Loomans *et al*., 2018; Georgakopoulos *et al*., 2020; Huch and Koo, 2015). Pancreatic islet cell transcription is also central to the mechanisms that underlie various forms of diabetes (Guo *et al*., 2013; Servitja and Ferrer, 2004; Miguel-Escalada *et al*., 2019).

Cellular programming and differentiation results from an interplay of positive and negative transcriptional regulatory mechanisms (Crews and Pearson, 2009; Graf and Enver, 2009). Several DNA binding transcription factors are known to promote endocrine differentiation during pancreas development (Gradwohl *et al*., 2000; Nishimura *et al*., 2006; Naya, Stellrecht and Tsai, 1995; Ahlgren *et al*., 1998; Stoffers *et al*., 1997; Matsuoka *et al*., 2004). A more limited number of transcription factors have been shown to act as negative regulators of endocrine differentiation (Jensen *et al.*, 2000; Martin *et al*., 2008; Cebola *et al*., 2015; Rosado-Olivieri *et al*., 2019; Mamidi *et al*., 2018). Some lines of evidence have pointed to RE-1 silencing transcription factor (REST, also known as NRSF, for neural-restrictive silencing factor) as a negative regulator of pancreatic endocrine differentiation (van Arensbergen *et al*., 2010). REST is a suppressor of neuronal genes in non-neuronal cell types (Chong *et al*., 1995; Schoenherr and Anderson, 1995). It is known to bind a 21-bp DNA recognition sequence, and has two repressor domains that recruit co-repressor complexes (reviewed in (Ooi and Wood, 2007; Hwang and Zukin, 2018)). Consistent with its function to inhibit neuronal genes, REST is largely expressed in non-neuronal cell types. However, REST is not expressed in endocrine β cell lines, and several genes that are repressed by REST are active in islet cells (Atouf, Czernichow and Scharfmann, 1997; Martin *et al*., 2008; Martin *et al*., 2012). Furthermore, genome-wide studies in embryonic stem cells and non-neuronal cell types have shown direct binding of REST near β cell-enriched genes (Johnson *et al*., 2008; Mukherjee *et al*., 2016; van Arensbergen *et al*., 2010). REST binding sites in embryonic stem cells overlap with genomic regions that carry Polycomb repressed chromatin in FACS-purified multipotent progenitors of the early embryonic pancreas (van Arensbergen *et al*., 2010). Many of these genomic regions are β cell regulatory genes that are subsequently derepressed during endocrine differentiation, in parallel with the concomitant loss of REST expression (van Arensbergen *et al*., 2010). These correlations have suggested that REST could be an important negative regulator of the endocrine differentiation program of the developing pancreas. A recent report further emphasized this notion, showing that REST curtails PDX1-mediated activation of endocrine genes in exocrine cells (Elhanani *et al*., 2020).

Genetic loss of function studies, however, failed to support the notion that REST plays a central essential role in pancreatic endocrine differentiation. Cre/LoxP-based excision of *Rest* in pancreatic progenitors led to changes in the expression of some endocrine genes but did not affect the number of endocrine cells, suggesting it was not an essential modulator of endocrine differentiation (Martin *et al*., 2015). Another pancreas deletion study reported that REST tempers pancreatic tissue damage and prevents acinoductal metaplasia, but did not assess endocrine differentiation (Bray *et al*., 2020). These studies, however, used an allele that removes *Rest* exon 2 (Gao *et al*., 2011). Recent work using a gene trap that disrupts transcription from all *Rest* promoters revealed dramatic effects on embryonic neurogenesis that were not observed when targeting *Rest* exon 2 (Nechiporuk *et al*., 2016). The same study showed that excision of Rest exon 2 does not prevent translation of a C-terminal REST peptide that is able to bind DNA, recruit co-repressors, and repress targets genes (Nechiporuk *et al*., 2016). Existing data, therefore, warrant a need to explore the true impact of REST in pancreatic endocrine differentiation using alternative genetic tools.

We have now inactivated *Rest* in the embryonic pancreas using a conditional allele that led to a drastic increase in endocrine differentiation, proliferation, and cell mass. We perform for the first time ChIP-seq studies of REST binding sites in the developing pancreas, and define salient properties of REST-dependent pancreatic program. We also show that endocrinogenesis can still be induced by inactivation of *Rest* in early postnatal duct cells, while this effect is dampened in adult mice. We used chemical inhibitors to show that REST function is conserved in zebrafish, and can also derepress endocrine genes in human pancreas organoids. Our results, therefore, demonstrate an essential role of REST as a major negative regulator of pancreatic endocrine differentiation.

## RESULTS

### REST expression in pancreas is largely restricted to duct cells

The expression of REST in pancreatic cell types has been difficult to resolve unequivocally due to low expression levels and lack of robust antibodies (van Arensbergen *et al*., 2010; Martin *et al*., 2015). We found nuclear REST immunoreactivity in most non-endocrine epithelial and mesenchymal cells of the mouse E12.5 pancreas, whereas from E14.5 onwards it was largely restricted to duct cell clusters and absent from acinar and endocrine cell clusters (**Supplementary Figure 1A**). Purified duct cells from adult and E18.5 *Sox9*-eGFP transgenic mice (Gong *et al*., 2003) confirmed *Rest* mRNA expression in duct cells (**Supplementary Figure 1B-C**). Finally, single cell RNA-seq data (Tabula Muris *et al*., 2018) showed *Rest* mRNA in adult duct and non-epithelial cells, but not in acinar or endocrine cells (**Supplementary Figure 1D**). These results reinforce the notion that REST is expressed in embryonic ductal progenitors and adult pancreatic ductal cells but is not detected in endocrine cells, consistent with a potential function of REST as a negative regulator of pancreatic endocrine differentiation.

### REST inactivation in pancreatic progenitors induces *Neurog3*

Previous genetic studies concluded that genetic ablation of *Rest* in pancreatic multipotent progenitors has no impact on the formation of NEUROG3+ endocrine precursors or hormone-producing cells (Martin *et al*., 2015), although this was examined in a mouse model that creates a deletion of *Rest* exon 2, which produces a functional isoform that can still bind DNA and recruit co-repressors (Nechiporuk *et al*., 2016). We thus used an allele that enables conditional excision of exon 4, which encodes >75% of REST protein residues (Yamada *et al*., 2010). Breeding this line with a *Pdx1-Cre* transgene (Gu, Dubauskaite and Melton, 2002) enabled the excision of *Rest* and a severe depletion of REST protein in most embryonic pancreatic epithelial cells (hereafter referred to as *Rest^pKO^* mice) **Supplementary Figure 2A-C**).

Previous work showed that REST binds to pancreatic endocrine regulatory genes in mouse ES cells, and that during embryonic pancreas differentiation REST target genes lose Polycomb-repressed chromatin and undergo transcriptional activation (van Arensbergen *et al.*, 2010). To directly test if REST truly acts as a repressor of endocrine differentiation during pancreas development, we examined expression of the endocrine lineage determinant NEUROG3 in *Rest^pKO^* embryos. This showed a 3.0 ± 0.03 -fold increase of *Neurog3* mRNA in *Rest*^pKO^ vs. *Rest*^LSL^ E13.5 embryos (SEM, Student’s T test p < 0.01), and 1.7-fold increased NEUROG3 protein (p < 0.05) (**Figure 1A-B**). At E18.5, *Neurog3* mRNA was 10.7 ± 1.2 -fold higher in *Rest*^pKO^ embryos (p < 0.01) (**Figure 1A**), and there was a ~6-fold increase in NEUROG3^+^ cells normalized by the total number of CK19^+^ cells (p < 0.01) (**Figure 1C**). Therefore, REST inactivation in pancreatic progenitors led to markedly increased of NEUROG3+ cells.

**Figure 1:**
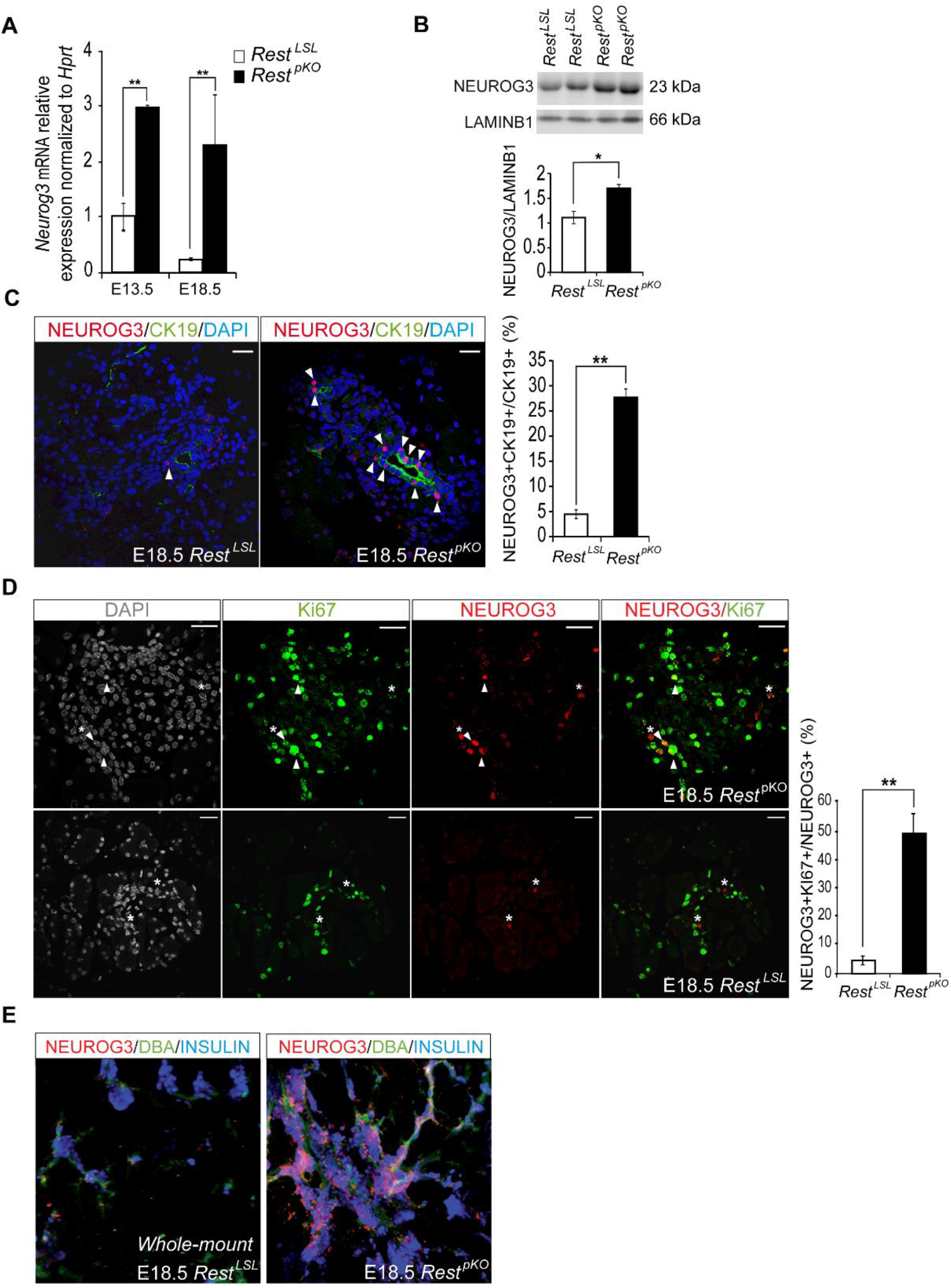
*Rest* inactivation in pancreatic progenitors induces NEUROG3. A) *Neurog3* mRNA increases in E13 and E18 *Rest*^pKO^ pancreas. Normalization by *Hprt* mRNA, n= 3-4 mice per group B) Western blot and quantifications of NEUROG3 in nuclear extracts from E13.5 *Rest^LSL^* and *Rest^pKO^* pancreas. LamininB1 was used as loading control. n= 2 samples per group with a pull of three E13.5 pancreas per sample. C) Immunofluorescence for NEUROG3 (red), Cytokeratin 19 (CK19, green) and DAPI (blue) in E18.5 *Rest*^LSL^ and *Rest^pKO^* pancreas. Arrowheads indicate NEUROG3+ cells. Bars show NEUROG3+ CK19+ cells in E18.5 pancreas. D) Immunofluorescence for NEUROG3 (red), Ki67 (green) and DAPI (grey) in E18.5 pancreas. Bars show NEUROG3+ Ki67+ cells. n=4-6 mice per group. E) Representative whole mounts for NEUROG3+ (red), DBA (green) and Insulin (blue) of the tail of E18.5 *Rest^LSL^* and *Rest^pKO^* pancreas. Scale bars = 100 μm, error bars are SEM. * p≤0.05 **p≤0.01.

The activation of *Neurog3* during normal pancreas development is followed by cell cycle exit of most progenitor cells (Miyatsuka *et al*., 2011; Krentz *et al*., 2017). We therefore investigated if REST inactivation affected the proliferation of NEUROG3-expressing cells. Immunofluorescence analysis of E18.5 *Rest^pKO^* pancreas showed that 48.3 ± 6.8 % of NEUROG3+ cells co-expressed Ki67 vs. 4.6 ± 1.5% in *Rest*^LSL^ control embryos (p < 0.01) (**Figure 1D**). This suggests that REST acts as a negative regulator of cell cycle exit in NEUROG3+ cells, which could contribute in part to the increased number of NEUROG3+ cells in *Rest^pKO^* embryonic pancreas.

NEUROG3+ cells arise from the pancreatic duct epithelium during a restricted developmental time frame (Gu, Dubauskaite and Melton, 2002; Solar *et al*., 2009). We tested if *Rest* deficiency could not only expand NEUROG3+ cells during embryonic development, but also cause persistent *Neurog3* activation throughout postnatal life. We therefore examined postnatal (2-week and 12-week-old) *Rest^pKO^* mice yet found no NEUROG3+ cells (**Supplementary Figure 3)**. This suggests that additional REST-independent mechanisms regulating *Neurog3* repression acquire prominence during postnatal life.

Our studies, therefore, show that REST tempers the expansion of NEUROG3+ cells in the embryonic pancreas, in part by suppressing NEUROG3+ cell proliferation, although it is not essential to prevent the continuous formation of endocrine progenitor cells from duct cells during postnatal life.

### Increased β cell mass in mice lacking REST in pancreatic progenitors

Given that the inactivation of REST in pancreatic progenitors led to the expansion of endocrine-committed NEUROG3+ progenitors, we investigated how this influenced the formation of endocrine cells. Whole-mount stainings for NEUROG3, CK19 and Insulin in E18.5 pancreas showed a marked increase of the number of insulin-expressing cells (**Figure 1E**).

Because the increase in NEUROG3+ cells observed in *Rest^pKO^* pancreas was transient, we investigated whether increased β cell mass was maintained in the adult pancreas. Young (12-16-week-old) male *Rest^pKO^* mice had normal weight (24.39 g ± 0.5 versus 24.36 g ± 0.6 *Rest^LSL^* control mice) and fasting glycemia (49.71 mg/dl ±1.83 versus 54.26 mg/dl ±3.44 *Rest^LSL^* control mice). Morphometric analysis, however, showed a ~2-fold increase of β cell mass in 12-week-old *Rest^pKO^* vs. control mice, which was primarily due to an increase of large islets (Student’s T test p < 0.01), as well as an apparent increase in glucagon-expressing cells (**Figure 2A-D**). These findings, therefore, showed that inactivation of *Rest* in pancreatic progenitors results in a transient expansion of NEUROG3+ cells and a sustained increase of pancreatic endocrine cell mass.

**Figure 2:**
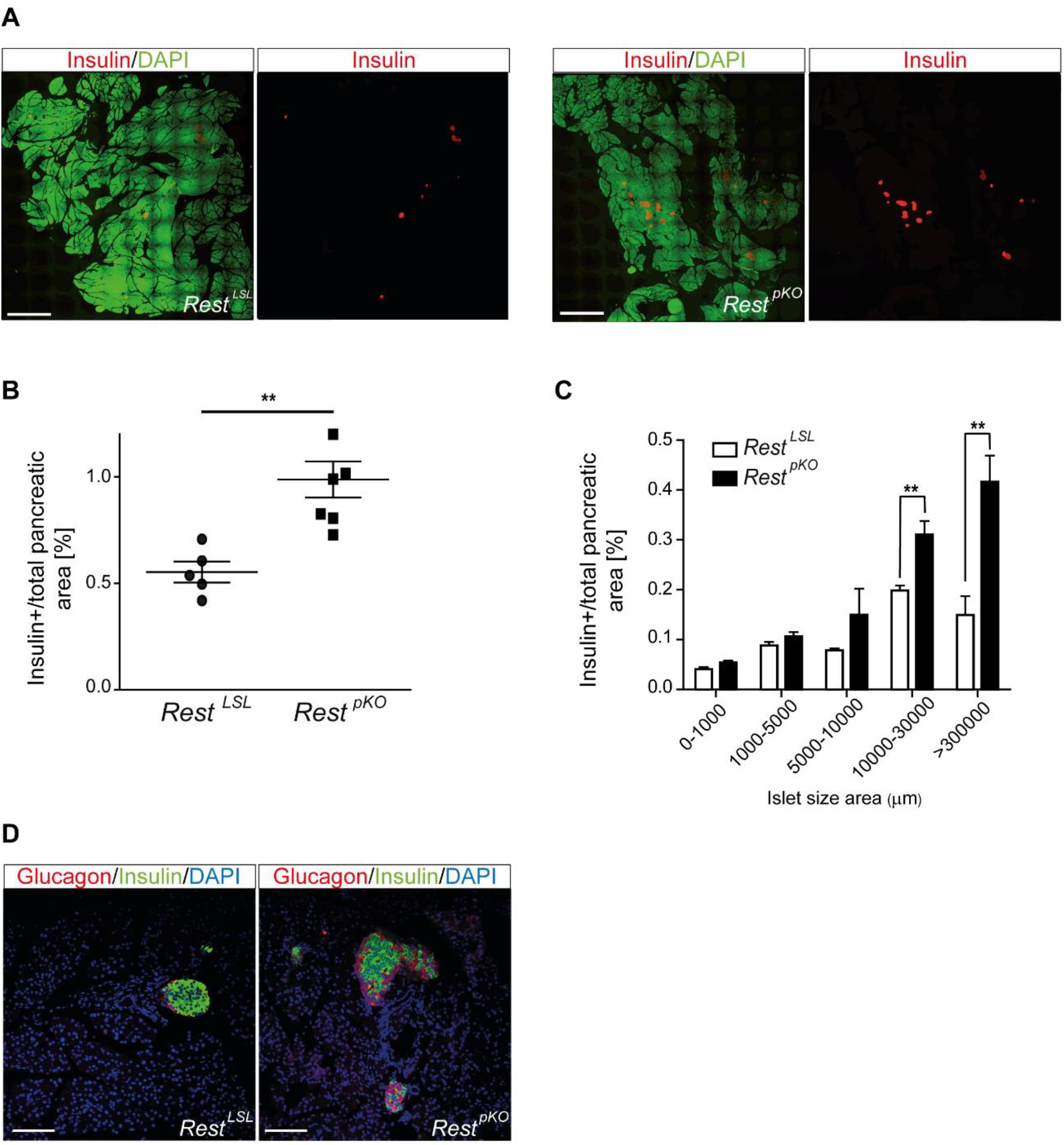
Increased β cell mass in *Resí*^pKO^ mice. A) Representative images of 10×10 frame reconstructions used for β cell morphometry of insulin (red) and DAPI (green) stainings in pancreas from 12 week old *Rest*^LSL^ and *Rest*^pKO^ mice. B) Morphometry of β cell mass estimated from insulin surface area/total DAPI surface area (%). Scale bar = 0.1 cm. *Rest*^pKO^ mice have ~2-fold increase β cell mass. n = 12 sections from 5-6 mice in each group. C) Islet size in *Rest*^LSL^ and *Rest*^pKO^ adult pancreas. D) Representative immunofluorescence for glucagon (red), insulin (green) and DAPI (blue) in 12-week-old *Rest*^LSL^ and *Rest*^pKO^ pancreas. Scale bar = 100 μm Error bars indicate SEM, **p≤0.01.

### REST is a direct regulator of pancreatic endocrine differentiation

Genome-wide REST binding maps from various cell types suggest that REST regulates different genes in different cell types (Hwang and Zukin, 2018), although REST binding sites have not yet been mapped in pancreas. To study how REST controls pancreatic endocrine differentiation, we identified REST genomic binding sites and REST-dependent transcriptional changes in embryonic pancreatic progenitors. We performed ChIP-seq analysis using a monoclonal antibody (12C11) directed to the REST C-terminal region (Chen, Paquette and Anderson, 1998), and used chromatin from E13.5 wild type pancreas –a stage in which REST expression is largely confined to progenitor cells and interspersed mesenchymal cells (**Supplementary Figure 1A**). We detected 1,968 REST-bound regions (**Supplementary Table 1**). These were highly enriched in canonical REST recognition motifs, confirming the specificity of REST binding (Otto *et al*., 2007) (**Supplementary Table 2** and **Figure 3A**). Integration with ATAC-seq profiles from E13.5 pancreas revealed that REST-bound regions were accessible to transposase cleavage, but had an accessibility footprint that was distinctly narrower than recognition sites of pancreatic activating transcription factors, which showed broader accessible regions, plausibly because active regulatory elements are bound by more regulatory proteins (**Supplementary Figure 4**).

**Figure 3:**
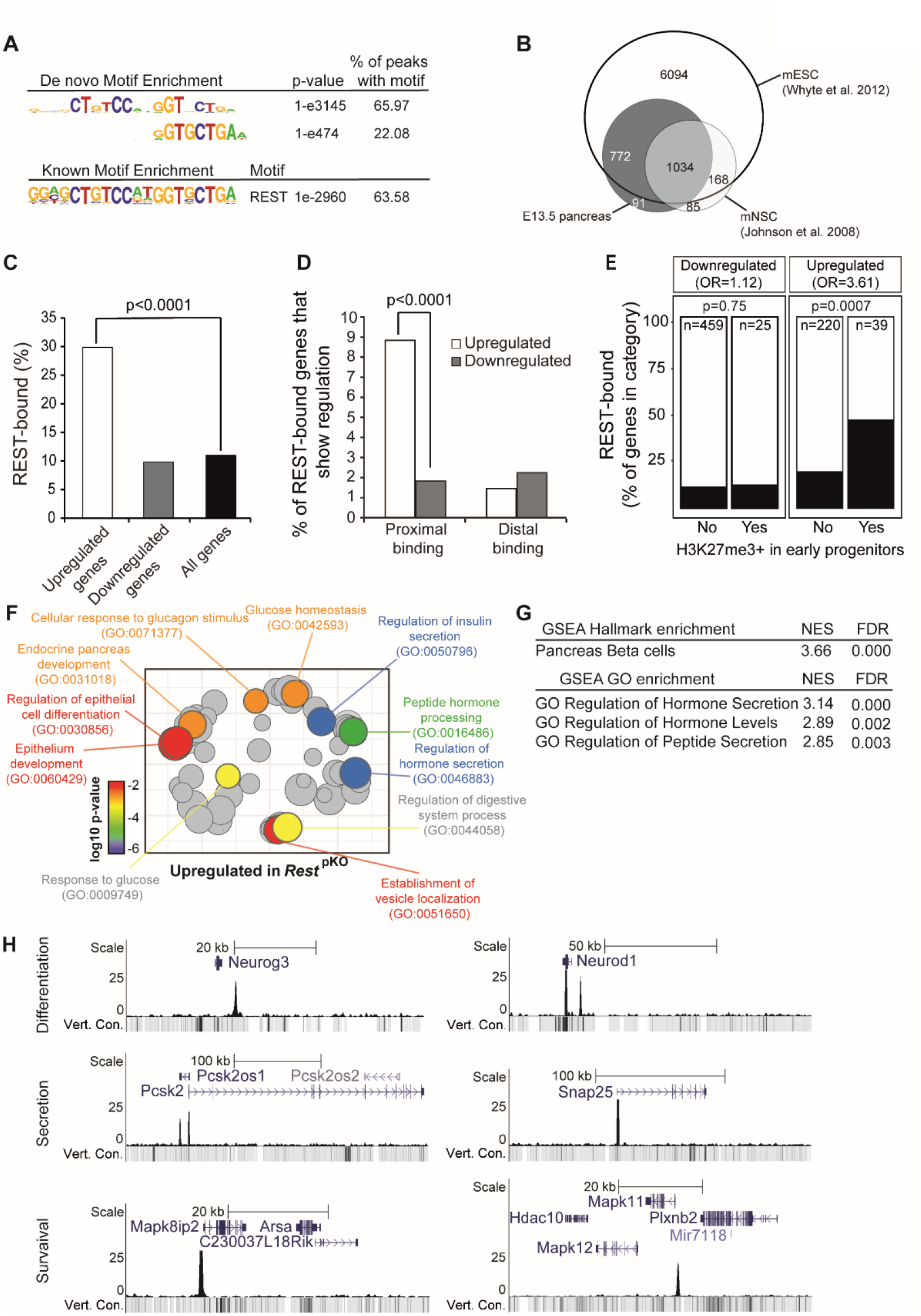
Functional direct REST targets in the embryonic pancreas. A) Top *de novo* and known motif enrichments in REST-bound regions. B) REST-bound regions in E13.5 pancreas, mESC and mNSC. C) Percentage of upregulated and downregulated genes in *Rest*^pKO^ mice, or all genes, that were bound by REST. Results indicate that REST predominantly acts as a repressor. p values are Fisher’s exact test. D) REST binds preferentially to promoter-proximal (0 - 5 kb) regions of genes that were upregulated in *Rest*^pKO^ mice. E) Percentage of differentially expressed genes that were bound by REST, broken down by H3K27me3 enrichment in purified pancreatic progenitors in (van Arensbergen *et al*., 2010). REST binding was enriched in genes that were upregulated in *Rest*^pKO^ and showed H3K27me3 in progenitors. P values from Fisher’s exact test. (F) Upregulated genes were functionally annotated using Gorilla (Eden *et al*., 2009), and REVIGO (Supek *et al*., 2011) was used to visualize annotation clusters. Most significant terms are highlighted according toa p-value colour scale. G) Significant GSEA terms for upregulated genes. H) REST binding associated to pancreatic endocrine development *(Neurog3 and NeuroD1),* insulin secretion *(Snap25, Pcsk2)* and β cell survival *(Mapk11* and *Mapk8ip2)* genes in E13.5 pancreas.

Many DNA binding transcription factors bind to different genomic regions in different cell types in which they are expressed (see for example (Servitja *et al*., 2009)), and this has been observed for REST (Hwang and Zukin, 2018). We found that 1,806 (~94%) of REST-bound sites in embryonic pancreas were shared with embryonic stem cells, but only 1,030 (~53%) with neuronal stem cells (Whyte *et al*., 2012; Johnson *et al*., 2008), while 91 (~4.7%) were exclusively detected in mouse embryonic pancreas (**Figure 3B**). These findings, therefore, defined direct REST-bound regions in early embryonic pancreas. They confirmed that numerous bound regions vary across cell types, although the vast majority are shared with embryonic stem cells (**Supplementary Table 3**).

To further assess REST function in pancreatic progenitors, we compared RNA-seq profiles of E18.5 *Rest*^pKO^ vs. *Rest*^LSL^ pancreas, a stage in which most progenitors have already been allocated to distinct cellular lineages. We identified 484 downregulated and 259 upregulated genes in *Rest^pKO^* E18.5 embryonic pancreas (adjusted p <0.05) (**Supplementary Table 4**). We found that 29.8% of upregulated genes were bound by REST (Fisher’s p < 10^-4^, relative to 15,548 expressed genes), whereas only 9.8% of downregulated genes were bound – a similar frequency as all expressed genes (11.2%; Fisher’s p = 0.81) (**Figure 3C and Supplementary Table 5**). This was consistent with the notion that the *Rest*^pKO^ phenotype reflects a transcriptional repressor function of REST in the developing pancreas.

Upregulated genes in *Rest*^pKO^ were selectively enriched amongst genes with promoter-proximal REST binding (Fisher’s p < 10^-4^) (**Figure 3D**). Furthermore, REST-bound upregulated genes were enriched in Polycomb-repressed chromatin in pancreatic progenitors (**Figure 3E**) (odds ratio=3.61, Fisher’ p = 7 x 10^-4^ for upregulated REST-bound genes; and odds ratio=1.12, p = 0.75 for downregulated REST-bound genes; both calculated relative to H3K27me3-enriched genes in PDX1+ E10.5 pancreatic progenitors) (van Arensbergen *et al*., 2010).

Consistent with the increase in endocrine cells of pancreas, genes that were upregulated, as well as pancreas REST-bound genes at large, showed a strong enrichment in pancreatic endocrine annotations, including insulin secretion and processing, glucose homeostasis, and endocrine pancreas development (**Figure 3F-G** and **Supplementary Table 6-7**). Closer inspection of individual loci disclosed the location of REST-bound regions at important regulators of pancreatic differentiation *(Neurog3, Neurod1, Insm1, Hnf4a, Onecut1, Pax4, Glis3, Hnf1a),* insulin biosynthesis or exocytosis *(Pcsk1, Pcsk2, Scg3, Snap25, Syt7),* as well as endocrine cell growth *(Bid, Mapk8IP1, Mapk10, Mapk11)* (**Figure 3H, Supplementary Table 3**). Regions bound by REST in embryonic pancreas but not in neural or embryonic stem cells included genes previously implicated in pancreatic differentiation and function such as *Isl1* (Ahlgren *et al*., 1997), *Ppdpf* (Jiang *et al*., 2008), *Fgf2* (Hebrok, Kim and Melton, 1998), *Shox2* (Cheng *et al*., 2019; Gu *et al*., 2004), *Prox1* (Paul *et al*., 2016), or *Cdh13* (Tyrberg *et al*., 2011) **(Supplementary Figure 5**). Given the increased proliferation of NEUROG3+ cells, it was also interesting to note upregulation of REST-bound positive cell cycle regulators in *Rest^pKO^,* including *Cdk5r2, Cdk2ap1, Ccnd1, Mapk3,* and *Ret* (**Supplementary Figure 5**, **Supplementary Table 5**). These studies, therefore, identified direct target genes through which REST controls pancreatic endocrine differentiation programs.

### Inactivation of REST in duct cells

Embryonic duct cells act as endocrine progenitors yet progressively lose their progenitor capacity (Solar *et al*., 2009; Kopp *et al*., 2011; Kopinke and Murtaugh, 2010; Kopinke *et al*., 2011), while REST expression is maintained as embryonic duct progenitor cells transition to adult differentiated ductal cells (**Supplementary Figure 1**). We therefore asked if REST inactivation immediately after birth could restore the capacity of duct cells to differentiate into endocrine cells. To this end, we used the *Hnf1b-CreERT2* transgenic line (Solar *et al*., 2009) and a Rosa26-LSL-RFP reporter (Luche *et al*., 2007) to trace ductal cells that have undergone *Rest* excision (**Figure 4A**). We treated dams of triple transgenic new-borns (hereafter called *Rest*^dKO^ Rosa26^RFP^) with tamoxifen at postnatal days 1 and 3, and then analysed mice after weaning (**Figure 4B**). This showed that the number of Insulin+/RFP+ and Glucagon+/RFP+ cells was increased 5.3±0.5 and 6.4±1.6-fold (Student’s T test p < 0.01), respectively, in *Rest*^dKO^ mice relative to *Hnf1b*-CreERT2 Rosa26^RFP^ control mice (**Figure 4C**). These results indicate that although the inactivation of REST in embryonic pancreatic progenitors does not result in persistent endocrine neogenesis throughout adult life, induced inactivation of REST in neonatal duct cells does allow them to transiently regain a progenitor-like state.

**Figure 4:**
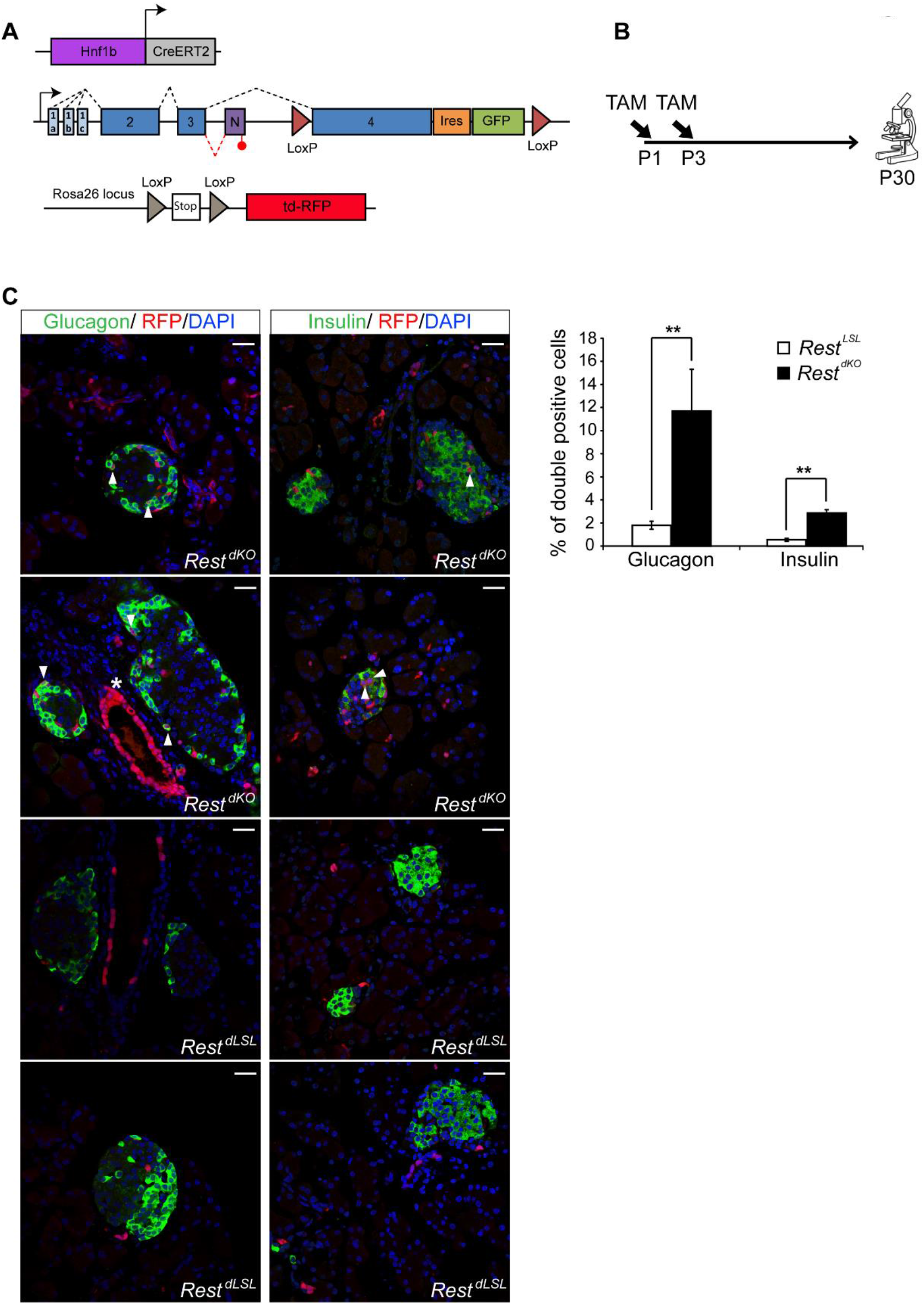
Pancreas-specific inactivation of Rest in neonatal ducts. A) Schematic of genetic models used to inactivate *Rest* and activate RFP expression in duct cells and progeny. *Hnf1b-CreERT2* is a BAC transgenic that specifically marks duct cells (Solar *et al*., 2009) and a small number of non-*Rest* expressing *∂* cells in reporters that are excised with high efficiency (Miguel Angel Maestro, Meritxell Rovira and Jorge Ferrer, unpublished observations), but not other endocrine or acinar cells. B) Schematic of the lineage tracing experiment. Tamoxifen was given to mothers at day 1 (P1) and day 3 (P3) after delivery, and mice were analysed at P30. *Hnf1b*-CreERT2Rosa26^RFP^ control mice were also treated. C) Representative images of double-positive RFP (red) and insulin (green) cells, and double-positive RFP (red) and glucagon (green) cells in *Rest*^dKO^ and control mice. The graph shows RFP-expressing glucagon and insulin cells in *Rest*^dKO^ vs. control mice. n=5-6 mice per each group. Arrowheads indicate double-positive cells, and an asterisk marks examples of cells in a duct, which were very efficiently labelled. Scale bar = 100 μm. Error bars are SEM. ** Student’s t test p<0.01.

By contrast, the inactivation of REST in ductal cells from 12-week-old mice did not lead to significantly increased of Insulin+/RFP+ cells (2.54±0.33% in *Rest^dKO^* Rosa26^RFP^ vs 1.96±0.49% in control mice (Student’s T test, p = 0.177) (**Supplementary Figure 6**). Thus, although REST retains an essential function to suppress endocrine cell formation from ducts in early postnatal periods, this role subsides in adult mice.

### Chemical inhibition of REST in zebrafish

Encouraged by the observation that inactivation of REST in early postnatal duct cells increased cellular multipotency, we explored if similar effects could be extended to other model systems using chemical inhibition of REST. We used X5050, recently identified in a high throughput screen to inhibit REST by protein destabilization (Charbord *et al*., 2013).

To study REST function in zebrafish, we used a double transgenic line where glucagon and insulin-expressing cells show green and red fluorescence, respectively *(Ins:mcherry/gcg:gfp).* We treated zebrafish embryos at 3 dpf with X5050, or with the Notch inhibitor DAPT as a positive control (Parsons *et al*., 2009), for 3 days. At 6 dpf we dissected the pancreas and quantified secondary islet formation as a read-out for endocrine cell differentiation. Compared to DMSO controls, the X5050-treated embryos displayed a dose-dependent increase in secondary islet formation (5 μM and 50 μM: 2.2±0.4 and 3.5±0.1-fold increase; SD, Student’s T test p < 0.05 and 0.01, respectively), comparable to 50 μM DAPT (4.2±0.4-fold, p < 0.01) (**Figure 5A**). These results indicate that REST regulation of pancreatic endocrine differentiation is conserved in zebrafish, and show that this process can be manipulated through chemical inhibition.

**Figure 5:**
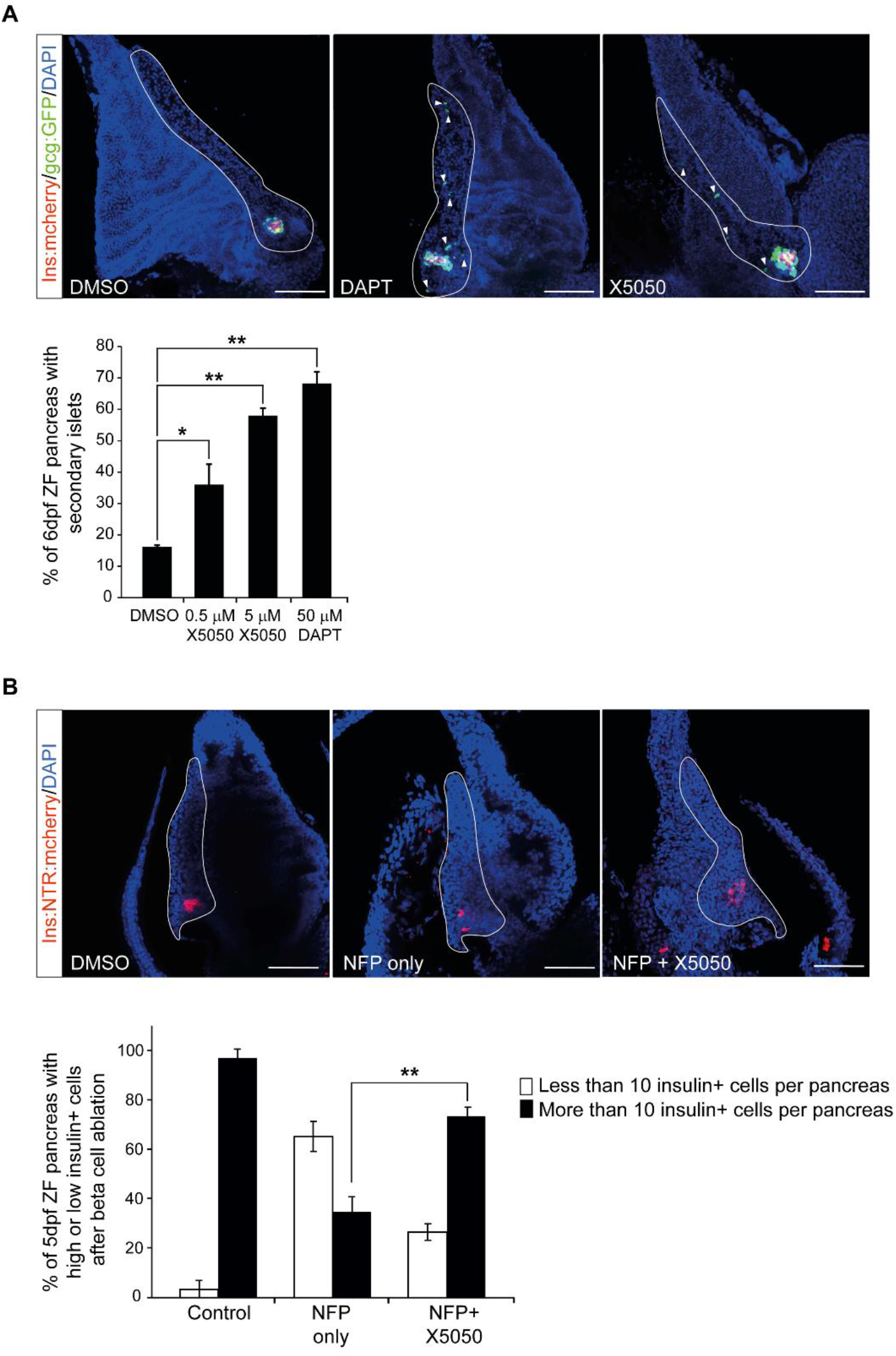
REST regulation of endocrine differentiation is conserved in zebrafish. A) Embryos of *Ins:mcherry/Gcg:GFP* double transgenic zebrafish line treated with DAPT (50 μM, Notch inhibitor used as positive control), X5050 (0.5 or 5 μM) or vehicle (DMSO, vehicle, negative control) from 3 dpf until 6 dpf. After drug treatment at 6 dpf zebrafish pancreas was dissected and presence of absence of secondary islets was quantified. Arrows show representative secondary islets of double transgenic zebrafish embryos *(Ins:mcherry/Gcg:GFP,* DAPI: Blue). The graph shows the percentage of zebrafish with detectable secondary islets in each condition. n= 20-25 fish per each condition. B) An *Ins:NTR:mcherry* line was used to selectively ablate β cells upon nifurpirinol (NFP) treatment (5 μM) (Bergemann *et al*., 2018; Pisharath *et al*., 2007) in 3 dpf embryos, and 24 hours later after complete β cell ablation of the principal islet, embryos were exposed to X5050 (5 μM) or vehicle, and β cells were analysed 36 hours later. Representative images of β cell regeneration in *Ins:NTR:mcherry* embryos treated with vehicle (DMSO), with NFP only, or with NFP and X5050. Insulin (red) and DAPI (blue). The graph shows the percentage of pancreas showing >10 insulin-expressing cells in every condition. n= 32-36 fish per each condition. Scale bars= 100 μm. Error bars are SEM ** p<0.01 and * p<0.05, Chi-square test.

We next investigated if REST inhibition could accelerate β cell neogenesis after β ablation in zebrafish. We used the *Ins:NTR:mcherry* transgenic line to selectively ablate β cells upon nifurpirinol (NFP) treatment (Bergemann *et al*., 2018; Pisharath *et al*., 2007) in 3 dpf embryos. In this model, 24 hours after NFP treatment >95% of β cells are ablated and β cell mass is recovered in 48-72 hours (Bergemann *et al*., 2018). We thus exposed embryos to NFP for 24 hours, and after washing NFP from β cell ablated embryos, these were treated with X5050 or vehicle and analysed for β cell recovery 36 hours later. We observed that β cell formation was accelerated upon REST inhibition, as 73.4 ± 3.3% of X5050-treated embryos showed >10 insulin-positive cells in the principal islet, while in control embryos only 34.7 ± 6.1% were observed (mean and SD of 34-38 embryos in each group, Chi-square test p < 0.01) **(Figure 5B).** Thus, REST inhibition promoted β cell formation in a zebrafish embryo regeneration model.

### REST inhibition in human organoids

We next explored the impact of REST manipulation in human cells. We validated the effect of X5050 in human cells using a human duct cell line, PANC-1, and showed that 24-hour treatment with X5050 resulted in a ~50% reduction of REST full length protein, and a relative increased in the REST4 isoform, as previously described (Charbord *et al*., 2013) (**Figure 6A**). Next, we studied *ex vivo* organoid cultures from human adult ducts isolated from the exocrine fraction of the pancreas of cadaveric donors. Organoids were generated and expanded as described in (Boj *et al*., 2015), and experiments were carried out at passage 3-4 (**Figure 6B**). Currently, the efficiency of endocrine differentiation from published *ex vivo* pancreas organoid protocols is very limited (Huch *et al*., 2013; Boj *et al*., 2015). We thus investigated if REST inhibition in pancreatic human organoids could promote the activation of pancreatic endocrine lineage genes as observed in mouse ablation studies. We treated human organoids with X5050 for 48 hours and observed very limited endocrine cells in treated and nontreated organoids. However, X5050-treated organoids showed induction of *INS*, *NEUROG3* and *PDX1* mRNA levels (2.27±0.43, 3.18±1.06 and 2.18±0.54-fold increased vs. DMSO, respectively; SD, Student’s T test p < 0.01), while the duct cell marker *SOX9* mRNA did not change (**Figure 6B**). These results, therefore, show that chemical interference of REST in adult human pancreas organoids did not lead to β cell formation, consistent with genetic findings in adult mice, although it induced the transcription of endocrine genes.

**Figure 6:**
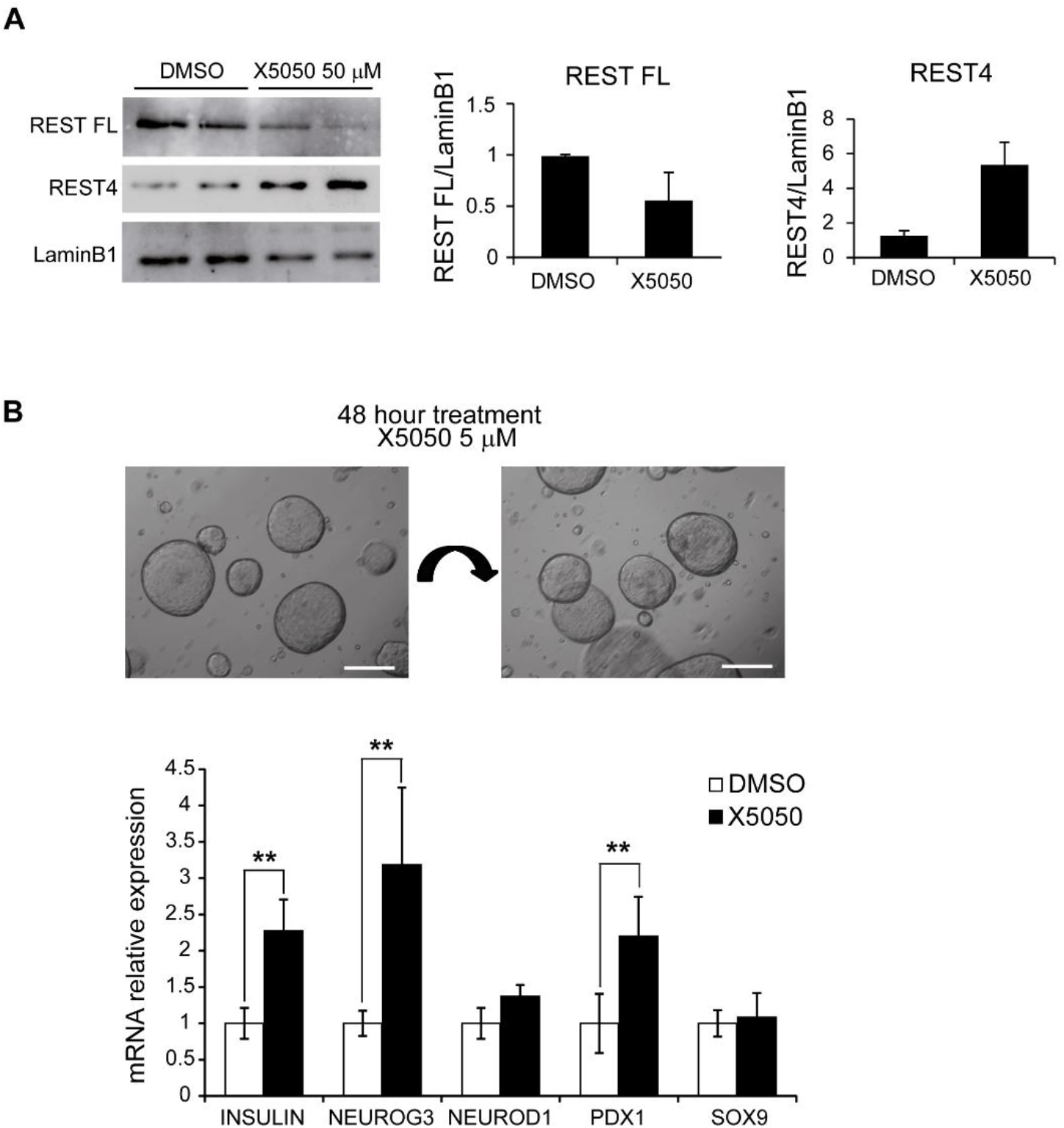
REST chemical inhibition in human pancreatic organoids. A) Western blot analysis of REST FL (fulllength) and REST4 protein levels in PANC1 cells treated with X5050 50 μM or DMSO (control). Lamin B1, loading control. Bar plot shows the quantification of the western blot for REST. B) Human organoids generated from pancreatic exocrine fractions from two cadaveric donors were treated at passage 2 for 48 hours with 5 μM X5050 or DMSO (control vehicle). qPCR analysis of mRNA for indicated genes, relative to TBP. Scale bars= 200 μm. Error bars are SD. ** Student’s t test p<0.01.

## DISCUSSION

Despite early suggestions that REST could be important for pancreatic endocrine differentiation during embryonic development (van Arensbergen *et al*., 2010; Martin *et al*., 2012; Martin *et al*., 2008), conditional ablation of *Rest* in the mouse pancreas unexpectedly showed modest gene expression differences and no quantitative changes in endocrine cell formation (Martin *et al*., 2015). This suggested that REST does not play an important regulatory role in pancreas endocrinogenesis. We have now used a different conditional allele that is known to cause more extensive ablation of REST function, and a Cre line that excises in embryonic pancreatic multipotent progenitors, and observed a drastic increase in pancreatic endocrine progenitors and hormone-producing cells. We show that this function of REST is conserved in a zebrafish β cell regeneration model. We define for the first time a blueprint of direct REST target genes in the embryonic pancreas, which underpin this regulatory function. We further demonstrate that the capacity to increase endocrinogenesis upon REST inactivation is retained in early postnatal duct cells, but not adult cells, while REST inihibition in human adult pancreas organoids influenced expression of endocrine genes but did not trigger endocrine cell formation.

Our findings, therefore, demonstrate that despite previous mouse genetic studies, REST is an essential major regulator of endocrinogenesis during pancreas development, and support a potential role of REST modulation for regenerative medicine.

Our findings should be assessed against the backdrop of current knowledge of positive and negative transcriptional regulators of endocrine cell formation from pancreatic multipotent progenitors. In addition to the established role of positive regulators of endocrinogenesis such as PDX1, NKX6-1, NKX2-2, RFX6, NEUROG3 and GLIS3 (Naya, Stellrecht and Tsai, 1995; Stoffers *et al*., 1997; Ahlgren *et al*., 1998; Gradwohl *et al*., 2000; Matsuoka *et al*., 2004; Nishimura *et al*., 2006), other DNA binding factors exert a suppressive function. This includes hippo-responsive TEAD-YAP complexes, which form part of pancreatic multipotent progenitor enhancers (Cebola *et al*., 2015; Rosado-Olivieri *et al*., 2019; Mamidi *et al*., 2018), as well Notch-responsive transcriptional repressors such as HES1). Our findings now establish that the REST-dependent program suppresses endocrinogenesis through interactions with promoters and distal regions, many of which show Polycomb repressed chromatin in multipotent progenitors. This result should prompt further studies to define how diverse positive and negative transcriptional regulatory programs interplay to control the formation of pancreatic endocrine cells.

The potential application of REST inhibitors to enhance endocrinogenesis from human pancreatic organoids is presently limited because existing protocols largely recapitulate exocrine cell expansion. A related problem is that we currently do not understand why REST de-repression can elicit endocrinogenesis in embryonic progenitors and even early postnatal duct cells, as well as in zebrafish models, but is much less efficient in the adult differentiated pancreas. Nonetheless, the observation that REST inhibition caused increased expression of islet endocrine genes in human exocrine organoids, as well as the effects on transcription factor-mediated reprogramming of mouse adult exocrine cells (Elhanani *et al.*, 2020), suggest that REST modulators may form part of an arsenal for future efforts to remove restrictions that impede endocrinogenesis in experimental model systems or replacement therapies.

## MATERIAL AND METHODS

### Mouse models

All experiments were approved by the Institutional Animal Care Committee of the University of Barcelona. Mice with *Rest* exon 4 floxed allele *(Rest^LSL^)* (Yamada *et al*., 2010) were crossed to *Pdx1*-Cre transgenic line (Gu, Dubauskaite and Melton, 2002) or *Hnf1b*-CreRT2 (Solar *et al*., 2009) and Rosa26-LSL-RFP transgenic line (Luche *et al*., 2007). We also generated *Rest^LSL^* mice carrying a different *Pdx1-Cre* transgene (Hingorani *et al*., 2003) and confirmed increased endocrine cell mass in adult mice as well as increase NEUROG3 positive cells in E18.5. To induce recombination in triple transgenics *(Rest^LSL^; Hnf1b-CreRT2;* Rosa26-LSL-RFP or *Hnf1b*-CreRT2; Rosa26-LSL-RFP control mice) at birth, 20 mg of tamoxifen were administered by gavage to the mother at day 1 and day 3 post-delivery, this allowing postnatal transfer of tamoxifen to the pups through breastfeeding. Mice were sacrificed at 30 days of age. Oligonucleotides used for genotyping are available in **Table 8**.

### Dissociation and FACS analysis of pancreatic cells

Adult and E18.5 mouse pancreas from *Sox9*-eGFP transgenic mice (Gong *et al*., 2003) were harvested and digested in 1.4 mg/mL collagenase-P (Roche) at 37 °C for 20-30 min. Peripheral acinar-ductal units, depleted of endocrine islets, were prepared as described in (Wang *et al*., 2013). Following multiple washes with HBSS supplemented with 5% FBS, collagenase-digested pancreatic tissue was filtered through 600 μm and 100 μm poly-propylene mesh (BD). Peripheral acinar-ductal units were further dissociated for FACS analysis in diluted TrypLE (Invitrogen) and incubated at 37 °C for 5 min. Dispersed cells were filtered through 40μm poly-propylene mesh (BD). Dissociated pancreatic exocrine cells were then resuspended at 1·10^6^ cells per ml in HBSS supplemented with 0.5% FBS. Cell sorting was performed using a FACS-Aria II (Becton Dickinson). The sorting gate for Sox9:eGFP positive ductal cells was established by using a WT mouse pancreas sample as negative control. Cells were directly sorted in RNeasy lysis buffer (Qiagen) for RNA extraction.

### Nuclear extracts and western blots

For isolation of nuclear extracts, E13.5 pancreas or PANC1 cells were washed twice with cold PBS. The tissue was resuspended in hypotonic lysis buffer (10 mM Hepes, 10 mM KCl, 0.1 mM EDTA, 0.1 mM EGTA, 1mM DTT) with protease inhibitors and place on ice for 15 minutes, followed by addition of 0.05% NP40 and vortex to disrupt cell membranes and then centrifuged at 12.000g at 4°C for 1 min. Pellet was resuspended in hypertonic buffer (10mM HEPES, 500mM NaCl, 0.1mM EDTA, 0.1mM EGTA, 1 mM DTT) with protease inhibitors and place on ice for 15 minutes with intermittent mixing by vortex. Centrifuge for 10min at 11.000g at 4°C. Supernatant containing nuclear extracts was kept in aliquots at −80°C.

Nuclear extracts were separated on a 7% SDS-PAGE gel and transferred to Immobilon polyvinylidene difluoride membrane (Millipore). The membranes were blocked with Tris-buffered saline Tween [10 mM Tris (pH 8.0), 150 mM NaCl, and 0.05% Tween 20] plus 5% BSA before incubation with the primary antibody (1:1000 for mouse monoclonal 12C11 anti-REST and 1:2000 for anti-LaminB1 rabbit polyclonal, Cell Signaling). Anti-rabbit IgG or anti-mouse IgG conjugated to horseradish peroxidase (1:2000) was used to detect the primary antibody. The membrane was washed with Tris-buffered saline Tween and the signal visualized using the Supersignal ECL reagent (Roche, Indianapolis, IN). Quantification of western blot signal was performed by using ImageJ.

### RNA analysis

Total RNA was isolated using the RNeasy Minikit (Qiagen) or Trizol followed by DNAse I treatment (Invitrogen). RNA was reverse transcribed with SuperScript III Reverse Transcriptase (Roche) and random hexamers according to the manufacturer’s instructions. qPCR of reverse-transcribed RNA samples was performed on a 7900 Real-Time PCR system (Applied Biosystems) using the Power SYBR Green reagent (Applied Biosystems). Quantities were determined using standard curves. Oligonucleotides are shown in **Table 8**.

### RNA-seq

All samples used for RNA-seq had RNA integrity number (RIN) > 8. RNA-seq data was generated from DNase-treated RNA from 3 independent E18.5 pancreases for each genotype (*Rest*^LSL^ and *Rest^pKO^)* using stranded Illumina library as 100 bp paired-end reads. Reads were aligned to the NCBI36/mm9 genome using star (v2.3.0) with default parameters, allowing only uniquely mapped reads. The resulting bam files were used to quantify gene expression using FeatureCounts (v1.5) using UCSC mm9 reference gene annotations. Differential expression analysis was performed using DESeq2 (Love, Huber and Anders, 2014) and genes that showed an adjusted p value below 0.05 were considered significantly differentially expressed.

### ChIP

ChIPs were performed as described in (van Arensbergen *et al*., 2010). In short, dissected E13.5 pancreatic buds were minced and rinsed with PBS, incubated for 10 min in 1% formaldehyde and 5 min in 125 mM glycine, rinsed in PBS containing protease inhibitor cocktail (Roche) at 4 °C, and snap-frozen and stored at −80 °C. Samples were sonicated using a Bioruptor (Diagenode) to achieve a chromatin size distribution of 200-1000 bp. Sonicated chromatin was diluted with ChIP dilution buffer (0.75% Triton X-100, 140 mM NaCl, 0.1% sodium deoxycholate, 50 mM HEPES at pH 8.0, 1 mM EDTA, 1× protease inhibitor cocktail) to achieve a final SDS concentration of 0.2%, pre-cleared with A/G Sepharose beads (GE Healthcare) for 1 h, incubated overnight at 4 ^°^C with 1 μg of REST antibody (12C11), rotated for 2 h at 4 ^°^C with A/G Sepharose beads, and then sequentially washed and processed. Immunocomplexes were eluted prior to DNA purification with Qiaquick columns (Qiagen).

### ChIP-seq

Chromatin from 3 independent pools of E13.5 pancreas were used for REST ChIP-seq experiments. All libraries were prepared using 1-2 ng of ChIP DNA and sequenced as single-end 50 bp reads using the Illumina HiSeq 2000 platform. Reads were aligned to NCBI36/mm9 genome using Bowtie2 (v2.2.5) allowing for one mismatch. The bam files were filtered to retain reads with a minimum mapping quality of 10 (MAPQ>=10) using samtools. The resulting bam files from biological replicates were pooled using samtools and peaks were called using MACS2 (v2.1.0) using default parameters. Bigwig files for visualization were generated using bamCoverage from deeptools2 using options – extReads 150 and –normalizeTo1x 2150570000.

### Functional annotation analysis

GSEA was performed in pre-ranked analyses with 1000 permutations, minimum term size of 10, and maximum term size of 500, using the stand-alone GSEA tool. Differentially expressed genes were functionally annotated using Gorilla (Eden *et al*., 2009), and REVIGO (Supek *et al*., 2011) was used to visualize annotation clusters with the following settings (0.9 allowed similarity; term size database-whole UniProt; semantic similarity measure-normalized Resnik; cluster definition default parameters) taking the most significant term in each GO cluster.

### ATAC-seq profiles around transcription factor binding sites (TFBS)

To plot ATAC-seq profiles around TFBSs of interest, we first conducted a footprinting analysis from the generated ATAC-seq data. For this purpose, we used the HINT framework from RGT library (v0.13.1) (Li *et al*., 2019). An ATAC-seq model and paired-end data were specified for tool execution. We considered the corresponding aligned read BAM file and the identified ATAC-seq peaks, in BED format, as the regions of interest. Required genomic data for mm10 assembly was obtained by running *setupGenomicData.py* script, available in the RGT suite. Once footprints were called, overlapping TFBSs were found using the matching motif analysis tool from RGT suite. By default, the complete JASPAR motifs database was used (Fornes *et al*., 2020). In the case of REST TFBS (MA0138.2), we selected only those footprinting regions that overlapped or were adjacent located to a REST ChIP-seq peak by using ChIPpeakAnno (v3.18.1) (Zhu *et al*., 2010), GenomicRanges (v1.36.0) (Lawrence *et al*., 2013) and Bioconductor (v3.9.0) R (v3.6.3) packages. Since REST ChIP-seq data referred to mm9 genome assembly, identified peaks were lifted over to mm10. Finally, normalized ATAC-seq profiles around the TFBSs of interest were generated with HINT for a 500bp window size. ATAC-seq bias-correction was enabled.

### REST binding enrichment in differentially expressed genes

We computed the fractions of REST-bound upregulated genes, REST-bound downregulated genes and (McLean *et al*., 2010) REST binding in all expressed genes. Genes were associated to REST-bound regions with GREAT v2.0.2 (McLean *et al*., 2010), applying default settings (basal plus extension; significant by both binomial and hypergeometric tests).

To analyse REST binding to proximal and distal genomic regions of down- and upregulated genes, REST-bound regions were binned by absolute distance to transcriptional start sites of annotated genes. Proximal and distal REST-bound regions were defined as < or > 5 kb from transcriptional start sites, respectively.

To analyse differential gene expression as a function of H3K27me3-enrichment and REST binding, we analysed H3K27me3-enrichment from (van Arensbergen *et al*., 2010). Genes were annotated to the corresponding BED files with GREAT v3.0 applying default parameters to basal plus extension association rule. To assess the association between H3K27me3+ and REST binding, odds ratio (OR) was obtained for each group. Fisher’s exact test was used to test association significance (p < 0.05).

### Motif analysis

*De novo* and known motif discovery of REST bound regions was carried out with HOMER (Heinz *et al*., 2010), using a 500-bp window centered on the REST peak sequence.

### Immunolocalization methods

Adult and embryonic pancreas were collected at indicated times and processed for immunohistochemistry of paraffin-embedded pancreas staining as previously described in (Maestro *et al*., 2003). Antibodies used are shown in **Table 9**.

Whole-mount staining of E18.5 pancreas were performed as described in (Ahnfelt-Rønne *et al*., 2007) without TSA amplification.

For β cell mass measurements, pancreases of 3-month-old mice were dissected and embedded in paraffin. Paraffin sections of (4 μm) were obtained at 150 μm intervals throughout the organs. Approximately 21 to 24 sections were analysed from each pancreas. Immunofluorescence for DAPI and insulin was performed. Images were taken by automated capturing and reconstruction of 10×10 frames using Leica SP confocal. Insulin-positive and total tissue areas (measured by DAPI saturation) were determined by ImageJ.

To assess islet size distributions, the sections analysed to study β cell mass were used to calculate islet size distribution. Islet size was measured using an automated plugin of ImageJ. Results were divided into 5 islet size groups of area in microns (<1000, 1000-5000, 5000-10000, 10000-30000, >30000).

### Human Organoid Culture

Human exocrine tissue was obtained from the discarded fraction from human islet purifications from cadaveric organ donors (Kerr-Conte *et al*., 2010). This procedure had permission to use tissue for scientific research if they were insufficient for clinical transplantation following national regulations, ethical requirements and institutional approvals from University of Lille. Ethical approval for processing pancreatic samples from deidentified organ donors was granted by the Clinical Research Ethics Committee of Hospital Clinic de Barcelona (HCB/2014/0926 and HCB/2014/1151).

Tissues was minced and digested with collagenase II (5 mg/ml, GIBCO) in human complete medium (Kerr-Conte *et al*., 2010) at 37° C for 30min to 1hr. The material was further digested with TrypLE (GIBCO) for 5 min at 37C, embedded in GFR Matrigel, and cultured in human organoid expansion medium (Boj *et al*., 2015) (AdDMEM/F12 medium supplemented with HEPES (1x, Invitrogen), Glutamax (1x, Invitrogen), penicillin/streptomycin (1x, Invitrogen), B27 (1x, Invitrogen), Primocin (1 mg/ml, InvivoGen), N-acetyl-L-cysteine (1 mM, Sigma), Wnt3a-conditioned medium (50% v/v), RSPO1-conditioned medium (10% v/v), Noggin-conditioned medium (10% v/v), epidermal growth factor (EGF, 50 ng/ml, Peprotech), Gastrin (10 nM, Sigma), fibroblast growth factor 10 (FGF10, 100 ng/ml, Preprotech) and PGE2 (1 mM, Tocris). After 3 passages we treated organoids with X5050 (Calbiochem) in human complete media for 48 hours prior to RNA analysis.

### Zebrafish drug treatments

The zebrafish lines used in this study were obtained from Isabelle Mandfroid (University of Liege, Belgium): *Tg(gcga:GFP), Tg(T2KIns:hmgb1-eGFP)* and *Tg(ins:NTR-mCherry). For* REST inhibition experiments double transgenic embryos *Tg(gcga:GFP)/ Tg(T2KIns:hmgb1-eGFP)* were incubated in 50 μM DAPT (Tocris) as a positive control and 0.5 and 5 μM X5050 (Calbiochem) (Charbord *et al*., 2013)) in E3 from 3 dpf until 6 dpf (Parsons *et al*., 2009). Embryos incubated in 1% DMSO in E3 were used as negative control. All the embryos were incubated at 28°C in dark during this period. After drug treatment 6 dpf zebrafish were fixed overnight in 4% paraformaldehyde, pancreas was then dissected and after confocal imaging of the pancreas presence or absence of secondary islets was quantified.

*Tg(ins:NTR-mCherry) was used for β* cell ablation studies upon nifurpirinol (NFP) treatment (5 μM) (Bergemann *et al*., 2018) in 3 dpf embryos. In this model, 24 hours post NFP treatment >95% of the β cells have been ablated and β cell mass recovers in 48-72 hours (Bergemann *et al*., 2018; Parsons *et al*., 2009). NFP-treated embryos were washed and treated with X5050 (5 μM) or DMSO. β cell regeneration was analysed after 36 hours based on the number of beta cells (red) per pancreas. After drug treatment 5 dpf zebrafish were fixed overnight in 4% paraformaldehyde pancreas was dissected and after confocal imaging of the pancreas the number of insulin positive cells in the principal islet per pancreas was quantified.

### Zebrafish embryo dissection and imaging

For confocal microscopy imaging, transgenic embryos were fixed overnight in 4% paraformaldehyde. To micro-dissect, fixed embryos were placed in PBS on an agarose lined plate. Then using pulled capillaries as tools, first the yolk and then the whole foregut were pried away from the embryo. The pancreas was placed islet down on a cover slip and dried by removing all excess. This cover slip was then mounted onto a microscope slide. Water was introduced under the cover slip to rehydrate the sample.

Confocal Z-series stacks were acquired on a Leica SP5 confocal microscope. Maximum projections were obtained by LAS AF software. To count endocrine cells we used a double transgenic line *Tg(gcga:GFP)* and *Tg(T2KIns:hmgb1-eGFP),* where glucagon positive cells are green and insulin positive cells are red. Upon maximum projections of Z-series of the entire pancreas presence or absence of secondary islets was computed.

### Statistics

Statistical analyses were performed using either R or GraphPad Prism 6. Experiments were analysed for statistical significance using Fisher’s exact test or unpaired, twotailed Student’s t-test with data expressed as mean ± S.E.M. unless otherwise specified as standard deviation (SD). P values < 0.05 were considered significant. Statistical methods and thresholds used for ChIP-seq and differential RNA-seq expression are described in corresponding sections.

## Acknowledgements

This research was supported by from Ministerio de Ciencia, Innovación y Universidades (MCIU) (SAF2015-73226-JIN and RYC-2017-21950) and European Research Council (FP-PEOPLE-2007-2-3-COFUND) to M.R. and grants to J.F. from Ministerio de Ciencia, Innovación y Universidades (BFU2014-54284-R, RTI2018-095666-B-I00), Medical Research Council (MR/L02036X/1), Wellcome Trust (WT101033), European Research Council Advanced Grant (789055). Work in CRG was supported by the CERCA Programme, Generalitat de Catalunya and Centro de Excelencia Severo Ochoa (SEV-2015-0510). With the support of the Instituto de Salud Carlos III, through the grant to JL.M. (CA18/00045) and MCIU fellowship to M.M. (PTA2018-016371-I). We thank the University of Barcelona School of Medicine animal facility, Imperial College London Genomics Units. We thank Cristina García for zebrafish support and maintenance. We thank Nikolay Ninov (Center for Regenerative Therapies, Dresden) and Isabelle Manfroid (University of Liège) for generous sharing of the zebrafish lines, and Irene Miguel-Escalada, Anthony Beucher and Inês Cebola for critical comments on the manuscript and discussions.

## Author Contributions

M.R. and J.F. conceived and coordinated the study. M.R. performed mouse, organoid, zebrafish and computational studies. M.R., M.A.M. performed image analysis of mouse mutants. G.A, M.M. and JL.M performed computational analysis. V.G. maintained mouse colonies and performed morphometry studies. J.G-H. performed ChIP experiments and RNA isolation. J.K-R and F.P. provided the pancreatic exocrine fraction of human donors for organoid cultures. M.R. and J.F. wrote the manuscript with input from the remaining authors.

## Data availability

Accession numbers pending

