## Supplementary Figures for "REST is a major negative regulator of endocrine differentiation during pancreas organogenesis"

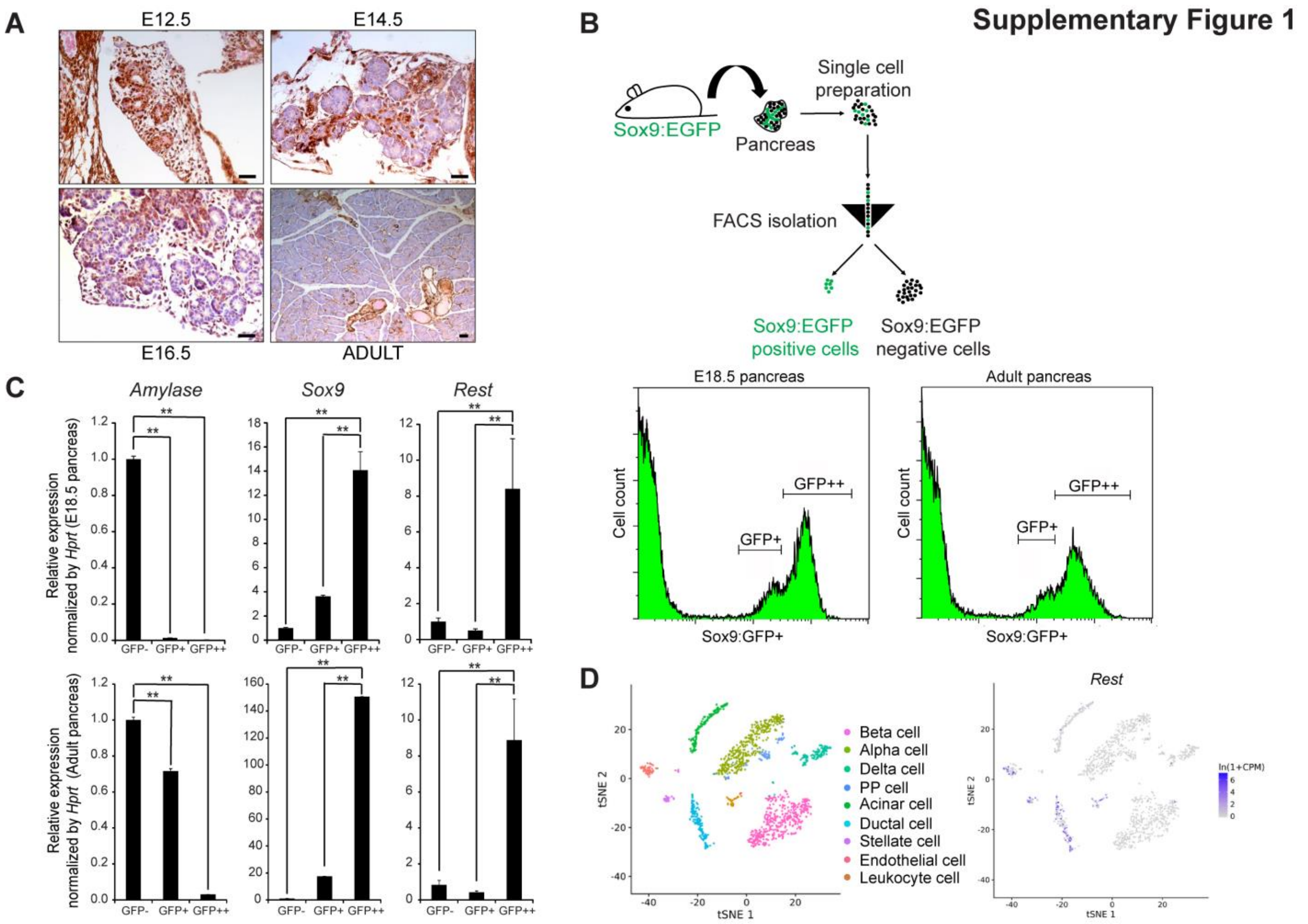


**Supplementary Figure 1: REST expression in the pancreas.** A) IHQ staining to detect REST protein levels during pancreas development (E12.6, E14.5 and E16.5) and adult. Scale bars= 100 µm. B) Schematic of the sorting strategy to isolate ductal cells from E18.5 and adult pancreas from *Sox9*-GFP transgenic mice (Gong *et al.*, 2003) . Representative sorting plots and gating strategy to isolate high (GFP++), low (GFP+) and negative (GFP-) Sox9-GFP expressing cells. C) qPCR analysis of *Rest* expression levels in FACS isolated ductal cells at E18.5 and adult pancreas, show increased *Rest* mRNA in duct-enriched (*Sox9* expressing cells) vs non-duct fractions (mainly acinar cells, *Amylase* expressing cells). D) Analysis of scRNA-seq datasets (Tabula Muris *et al.*, 2018), show *Re*st expression in adult pancreatic ductal cells and non-epithelial cells, but not in insulin- and glucagon-expressing cells. Error bars = ± SEM. ** p≤ 0.01.

### Supplementary Figure 2

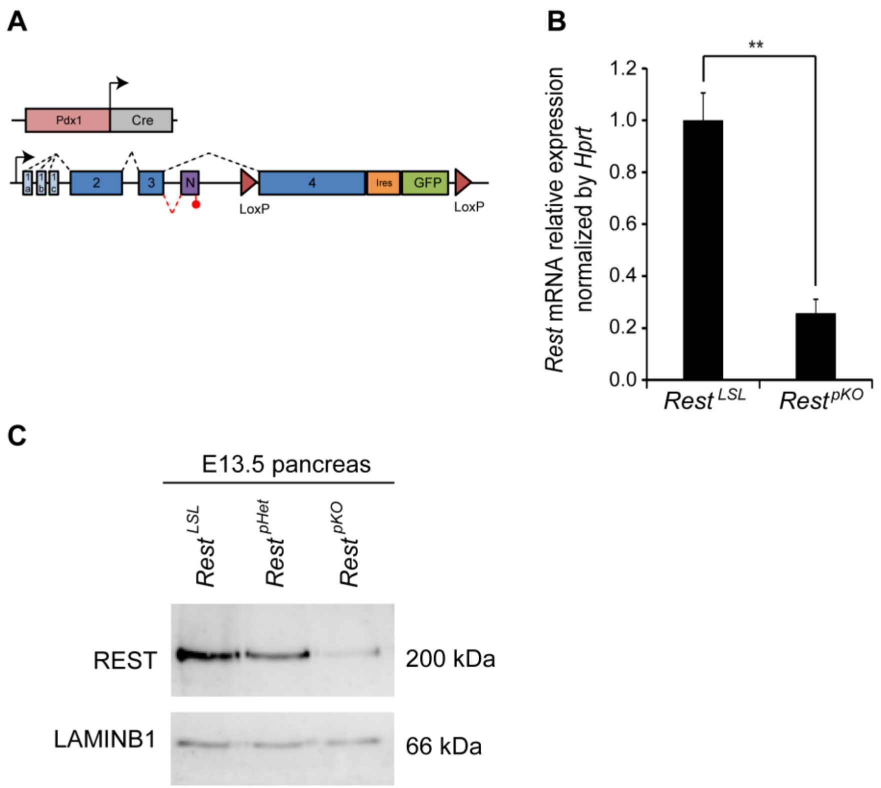


**Supplementary Figure 2: *Rest* pancreatic KO mouse model (*Rest*$^{pKO}$).** A) Schematic of the two genetic models to delete *Rest* from pancreatic epithelial cells. B) qPCR analysis of *Rest* mRNA levels in E13.5 pancreas upon deletion. Results are normalized by *Hprt*, n = 3 independent embryos in each group. Error bars are SEM. ** p≤ 0.01. C) Western blot images of REST protein levels in the pancreas of *Rest*$^{LSL}$ controls, *Rest*$^{pHet}$, carrying heterozygous LoxP allele and Pdx1-Cre, and *Rest*$^{pKO}$. LaminB1 is a loading control.

### Supplementary Figure 3

**A**

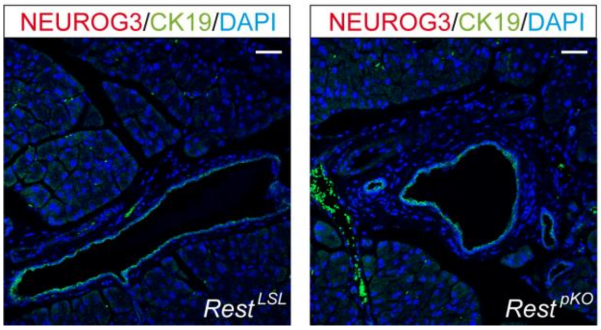


**Supplementary Figure 3: NEUROG3 expression in adult *Rest*$^{pKO}$ mice.** Representative immunofluorescence staining for *Neurog3* (red), CK19 (green) and DAPI (blue) in adult pancreas from *Rest*$^{LSL}$ *and Rest*$^{pKO}$ mice shows absence of NEUROG3 positive cells. These findings indicate that REST-independent inhibitory mechanisms, and or lack of *Neurog3* activating mechanisms, acquire prominence during postnatal life. Scale bars= 100 µm.

**A**

### Supplementary Figure 4

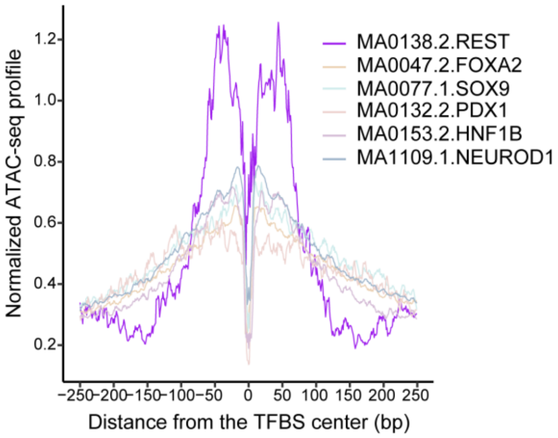


**Supplementary Figure 4:** Integration of REST-bound regions with ATAC-seq profiles from E13.5 pancreas shows a distinct chromatin accessibility footprint of REST-bound regions from activating transcription factors.

### Supplementary Figure 5

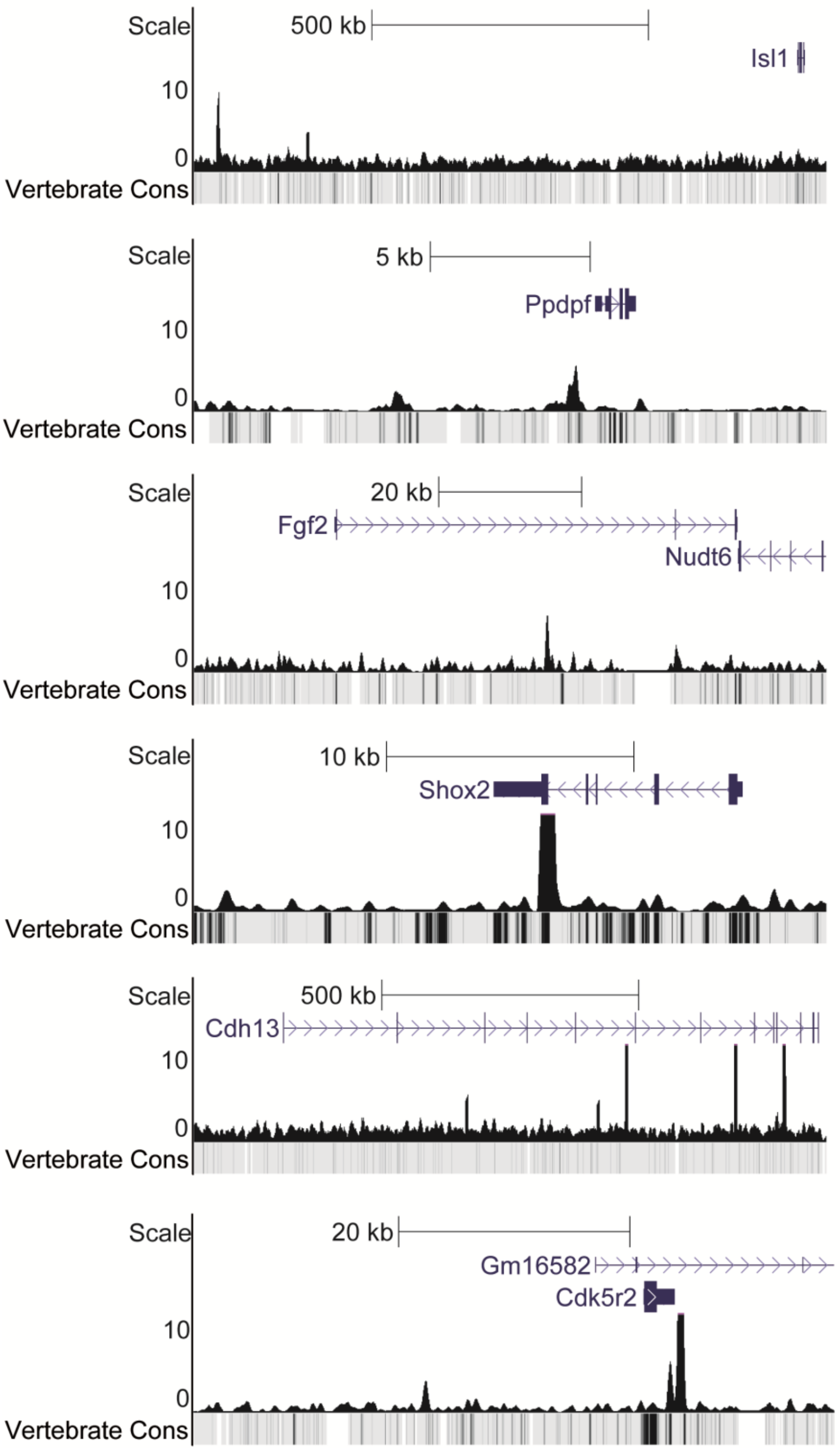


**Supplementary Figure 5: Representative tracks of uniquely bound REST-regions in embryonic pancreas versus mESC and mNSC**.

### Supplementary Figure 6

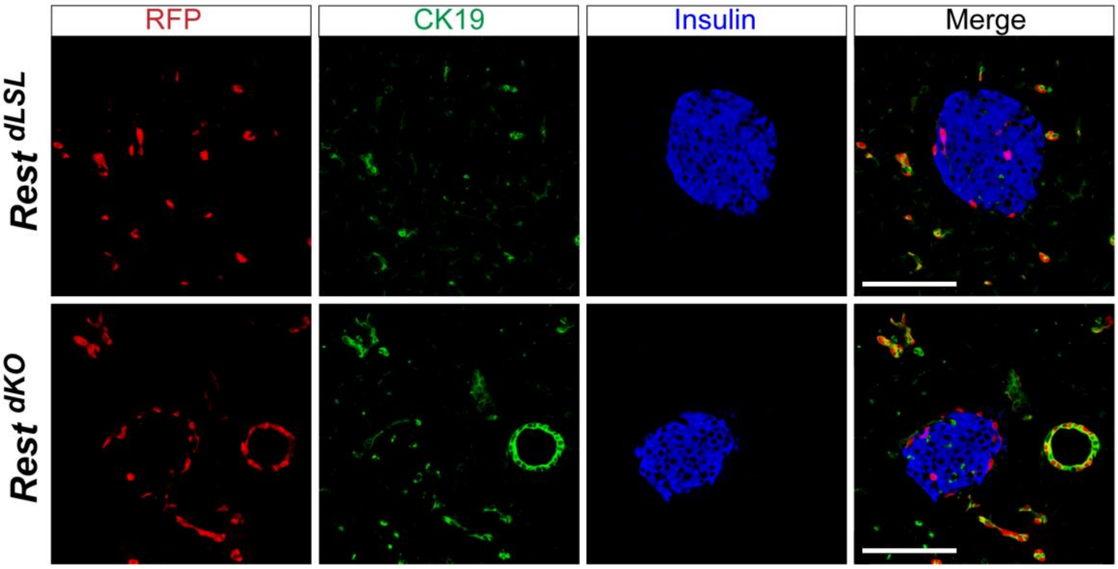


**Supplementary Figure 6: *Rest* pancreatic ductal KO in adult mice (*Rest*$^{\mathbf{dKO}}$*)*.** Representative immunofluorescence staining for *Insulin* (blue), CK19 (green) and RFP (red) in adult pancreas of 12-week-old mice from *Rest*$^{LSL}$ *(Hnf1b*-CreERT2 and Rosa26$^{RFP}$) *and Rest*$^{dKO}$ (*Hnf1b*-CreERT2 and REST LSL and Rosa26$^{RFP}$ genotypes. No differences in Insulin+/RFP+ cells between *Rest*$^{LSL}$ *and Rest*$^{dKO}$ mice were observed. Scale bars= 100 µm.
